# Molecular details of protein condensates probed by microsecond-long atomistic simulations

**DOI:** 10.1101/2020.08.05.237008

**Authors:** Wenwei Zheng, Gregory L. Dignon, Xichen Xu, Roshan M. Regy, Nicolas L. Fawzi, Young C. Kim, Robert B. Best, Jeetain Mittal

**Author notes:** These authors contributed equally to this work (Wenwei Zheng, Gregory L. Dignon).

## Abstract

The formation of membraneless organelles in cells commonly occurs via liquid-liquid phase separation (LLPS), and is in many cases driven by multivalent interactions between intrinsically disordered proteins (IDPs). Molecular simulations can reveal the specific amino acid interactions driving LLPS, which is hard to obtain from experiment. Coarse-grained simulations have been used to directly observe the sequence determinants of phase separation but have limited spatial resolution, while all-atom simulations have yet to be applied to LLPS due to the challenges of large system sizes and long time scales relevant to phase separation. We present a novel multiscale computational framework by obtaining initial molecular configurations of a condensed protein-rich phase from equilibrium coarse-grained simulations, and back mapping to an all-atom representation. Using the specialized Anton 2 supercomputer, we resolve microscopic structural and dynamical details of protein condensates through microsecond-scale all-atom explicit-solvent simulations. We have studied two IDPs which phase separate *in vitro*: the low complexity domain of FUS and the N-terminal disordered domain of LAF-1. Using this approach, we explain the partitioning of ions between phases with low and high protein density, demonstrate that the proteins are remarkably dynamic within the condensed phase, identify the key residue-residue interaction modes stabilizing the dense phase, all while showing good agreement with experimental observations. Our approach is generally applicable to all-atom studies of other single and multi-component systems of proteins and nucleic acids involved in the formation of membraneless organelles.

## Introduction

Biomolecular condensates are highly concentrated subcellular assemblies of biomolecules that occur naturally in biology, and may function as organelles, such as the nucleolus,^1–3^ ribonucleoprotein granules,^4,5^ and many others.^6–10^ The study of these bodies, often termed membraneless organelles (MLOs), has recently attracted tremendous research effort due to its novelty, and relevance to biological functions,^11–14^ pathologies such as neurodegenerative diseases^15,16^ and the design of biomimetic materials.^17–19^ It is now accepted that many MLOs are formed through a process of phase separation, commonly liquid-liquid phase separation (LLPS) in which a dynamic liquid-like condensate organizes biomolecules including proteins and nucleic acids and allows them to diffuse freely within the condensate, and to exchange rapidly with the surrounding environment.^7^ A physical understanding of the driving forces of biomolecular phase separation is essential for uncovering the mechanistic details of MLO formation and the pathology of relevant diseases.^20–25^

A frequent property of proteins involved in biomolecular phase separation is intrinsic disorder, which has been highlighted through estimates of enhanced disorder predicted within MLO-associated proteins.^26^ Indeed, intrinsically disordered proteins (IDPs) have been shown to phase separate at relatively low concentrations compared to most folded proteins,^5,25,27^ likely due to their polymeric nature, and consequent increased multivalent interactions. Additionally, IDPs are generally more solvent exposed,^28^ and thus more accessible to post-translational modifications, which provide an efficient mechanism of controlling the thermodynamic and dynamic properties of condensates.^23,29,30^ Recent work has highlighted that single-molecule behavior of IDPs may yield information relevant to their phase behavior, since the intramolecular interactions driving single-chain collapse are related to the intermolecular interactions driving its homotypic phase separation.^31,32^ This leads to the question of what exactly are the interactions that drive LLPS, how can they be determined, and how can they be manipulated to control phase behavior?^25^

Despite the advances in methodology for investigating structure formation inside LLPS droplets by experiment,^22^ it is still challenging to obtain high resolution, sequence-resolved information on structure and dynamics from experiment alone. All-atom molecular dynamics simulation with explicit solvent is a promising technique for generating detailed information on conformational ensembles of IDPs,^33,34^ and the contacts occurring within a condensate composed of IDP molecules.^22,35,36^ The approach has already been applied to simulating the condensed phase of disordered peptides and proteins.^37–39^ However, the large system sizes and timescales required to observe equilibrium coexistence of two phases pose a major challenge for all-atom simulations. We have previously overcome this difficulty by developing coarse-grained (CG) simulation models for LLPS.^29,31,40–43^ We have complemented CG simulations of LLPS with atomistic simulations of smaller fragments of the IDPs,^5,22,24–35,44^ which yield detailed interactions occurring between the relevant proteins in dilute solution, and in principle, within the condensed phase.^31^ In this work, we unify these two approaches, by using CG simulations to generate an initial, equilibrated configuration of phase-separated proteins, which is then mapped back to all-atom coordinates to investigate the details of atomic interactions occurring within a protein condensate.

## Results and Discussion

As test systems, we have selected the low-complexity prion-like domain of FUS (hereafter, FUS LC),^4^ and the disordered N-terminal domain of LAF-1 (LAF-1 RGG).^35,45^To set up the system, we initially equilibrated 40 chains of FUS LC or LAF-1 RGG in a planar slab geometry using our previously developed CG model^31,40^ (Fig. 1a). The system size was chosen as it yields an atomic resolution system that is sufficiently small to run on Anton 2, while being sufficiently large that finite size effects are small. We verify this by comparing with a larger system as we have used previously with 100 chains^31,40^ and find similar coexistence densities (see supporting methods and Fig. S1). After setting up this system, we reconstructed all-atom coordinates on the C_*α*_-only configuration by using a lookup table based on the protein structure database with the PULCHRA code^46^ (Fig. 1b). Any conflicts between sidechains of different chains were resolved via a short simulation with the CAMPARI Monte Carlo engine and ABSINTH implicit solvent model with fixed backbone^47^ (Fig. 1c). Finally, the system was solvated and equilibrated with explicit solvent using the Amber ff03ws^48^ force field, TIP4P/2005 water model,^49^ and ∼100 mM NaCl^50^ (Fig. 1d). By utilizing the specialized software and hardware from Anton 2 supercomputer developed by DE Shaw Research,^51^ we equilibrated the system for 150 ns to relax it to its equilibrium density (Fig. S2) and collected a 2 *µ*s trajectory in the NVT ensemble at 298 K for each sequence of interest (see supporting methods for details).

**Figure 1.**
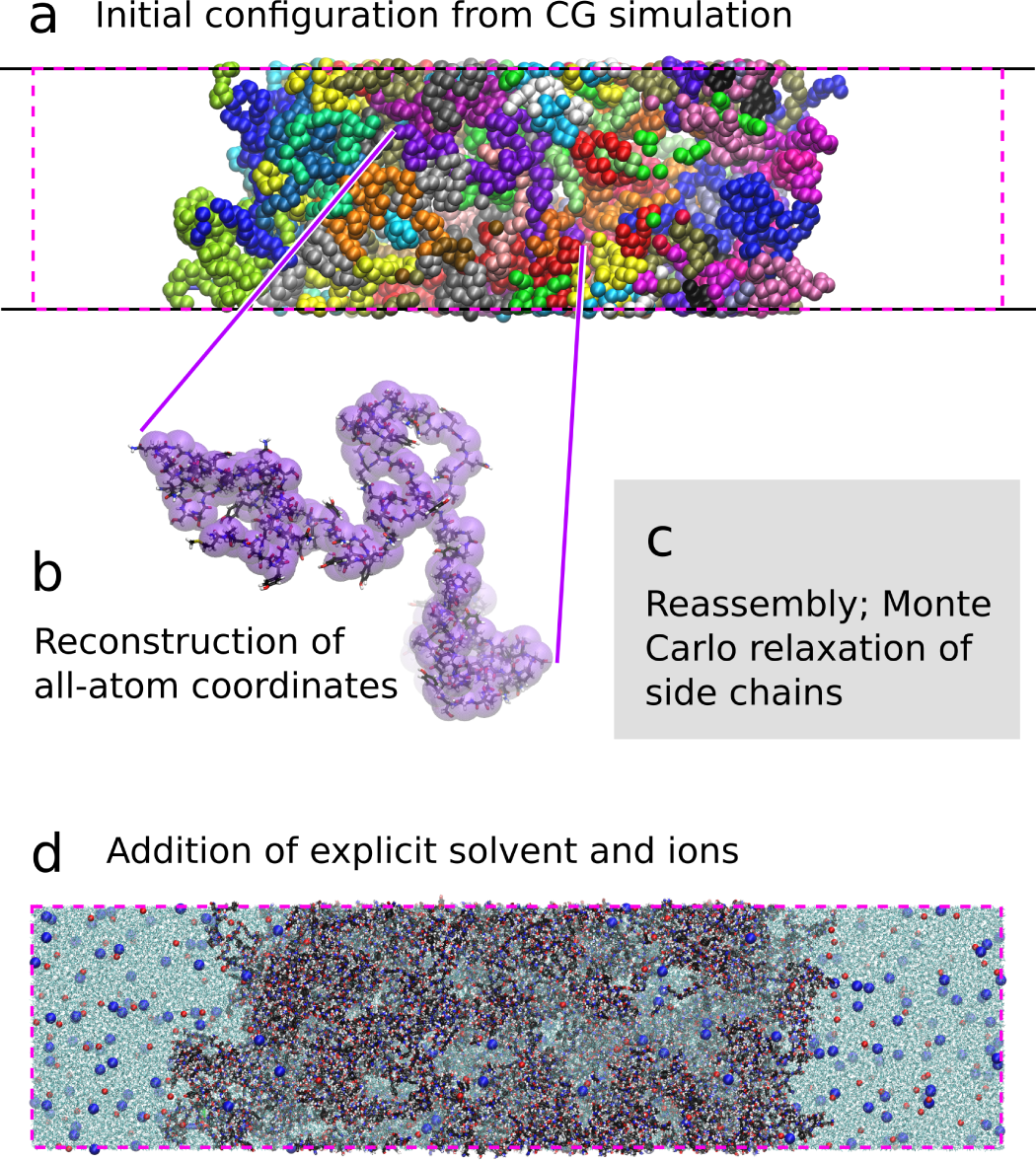
Procedure to set up all-atom simulations of dense phase. (a) An initial configuration is generated by CG simulations in a box with elongated *z*-dimension. (b) All-atom coordinates reconstructed from CG C_α_ coordinates templates for each chain using PDB database. (c) Reconstructed chains reassembled into condensed phase and sidechain clashes relieved using short Monte Carlo simulations with frozen backbone. (d) Addition of solvent and ions to generate complete system in all-atom resolution.

All-atom simulations with explicit solvent can provide a great deal of information not accessible from CG models, most obviously how the solvent and ions partition into the dense phase, and how this depends on protein sequence. The diffusion of solvent and ions is very rapid and so their equilibrium partitioning can be readily probed from our all-atom simulations, as shown in Fig. 2 for FUS LC and LAF-1 RGG protein condensates. The initial protein concentration in the dense phase for both proteins was selected based on the estimates from NMR experiments on FUS LC to be ∼477 mg/mL,^22^ and typical for protein LLPS.^52^ We note that there is also some indirect evidence for extremely low density condensates of LAF-1 RGG under certain conditions but it is not consistent with directly measured values for a human homolog ddx4 with very similar sequence which forms very high density phases.^52,53^ In both cases, the protein-rich phase has a higher total density than water (black lines), which agrees with the experimental observations that condensates of these proteins can be sedimented or separated using centrifugation.^4,45^ The water content inside both FUS LC and LAF-1 RGG protein-rich regions is on the order of ∼600 mg/mL (Figs. 2a and b), very similar despite significant differences in their sequence composition. The water content inside the FUS LC protein-rich phase from the simulation is consistent with the reported experimental estimate of 65% (by volume) by Murthy et al. ^22^

**Figure 2.**
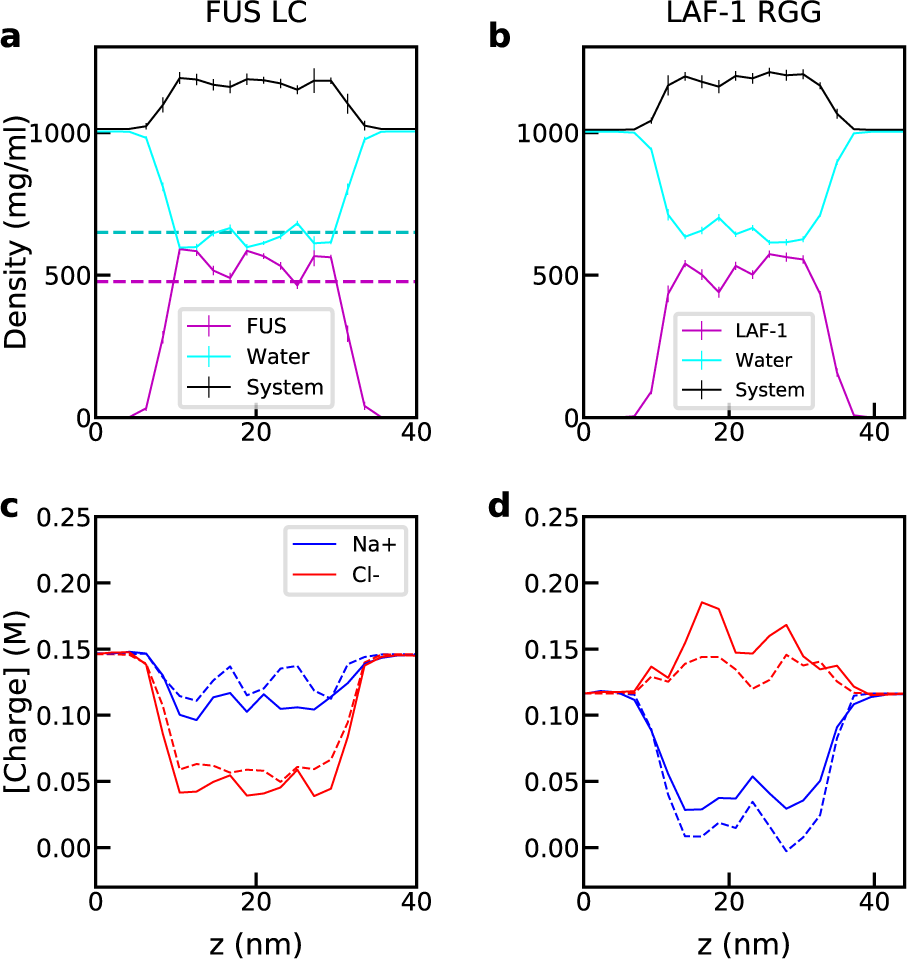
Density profiles from all-atom slab simulations of FUS LC (left) and LAF-1 RGG (right). Components are shown in the legend. Dashed lines in (a) indicate experimentally determined values^22^ and dashed lines in (c) and (d) indicate the predicted ion concentration using concentrations of protein cationic and anionic residues.

Despite very similar protein and water density profiles for FUS LC and LAF-1 RGG, the partitioning of Na^+^ and Cl^−^ ions differs considerably between the two systems (Fig. 2c and d). In the case of FUS LC, which only has two anionic residues (Asp), the concentration of Cl^−^ ions is greatly reduced inside the protein-rich phase, being preferentially excluded, while Na^+^ ion concentration is only slightly reduced in the protein-rich phase (Fig. 2c). For LAF-1 RGG (Fig. 2d), which contains a more significant fraction of anionic and cationic charged residues (26%), the Cl^−^ ions are preferentially incorporated into the protein-rich region, while the Na^+^ ions are excluded. This likely has to do with the net +4 charge per protein chain for LAF-1 RGG. The equilibrium partition coefficient of ions reflects an interplay of direct charge-charge interactions between charged amino acids and ions and the free energy of transferring the ions from a solvent-rich to a protein-rich environment. Using a simple model, we can predict the local concentration of Na^+^ and Cl^−^ from the local concentration of cationic and anionic residues (Fig. S3) and bulk concentrations of ions and water. We set

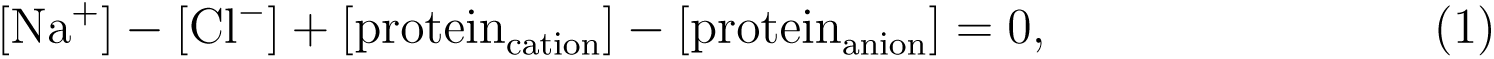

to represent electroneutrality, and

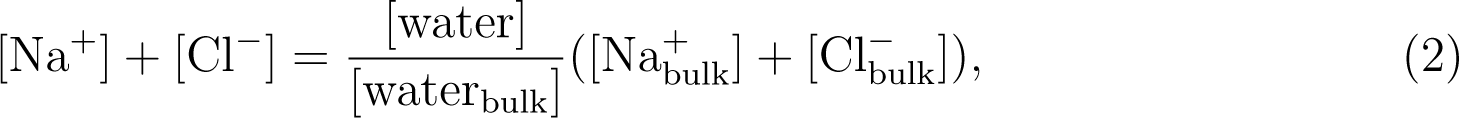

which assumes negligible preferential interactions between ions and amino acids, as would be expected at these relatively low ion concentrations. The predicted Na^+^ and Cl^−^ concentrations are plotted in Figs. 2c and d as dashed lines, and show good agreement with the concentrations obtained from the simulation. These results highlight the role of the charged amino acids in determining the density and composition of the protein condensates, which ultimately help to determine their function.

While equilibrium concentrations and compositions of MLOs are important to their function, another important factor is the dynamics within the dense phase, as it determines the rate at which components may pass through or rearrange within the condensate. Our MD simulations also provide detailed information on the dynamics within the condensed phase, and may be used to decouple the different components. The heterogeneous nature of our system with a distinct protein-rich environment, protein-poor bulk region, and an interfacial region poses some challenges to estimate diffusion coefficients unambiguously using the standard approach based on the mean square displacement. An alternative approach is to compute the probability distributions *P* (*ξ*(*t*_0_ + *t*) − *ξ*(*t*_0_)) for molecular displacement (i.e. propagators) in each direction *ξ* = *x, y, z* as a function of the lag time *t* between observations (see supporting methods and Fig. S4). Since the simulation box is not cubic, we report diffusivity (*D*) values based on only the longer *z*-axis, in order to minimize the finite size effects.^54^ We find it necessary to include more than one term (multiple *D* values) while fitting the propagator data from simulation to the expected distribution for one-dimensional diffusion. This behavior is consistent with the expected differences in the dynamics of solvent molecules within the protein-rich and bulk phases. We find that the observed behavior of water and ions is best accounted for by three *D* values whereas one *D* value is sufficient for fitting protein propagator data (Fig. S4).

The fastest *D* value for water (*D*_1_ ≈ 1.98 nm^2^/ns in FUS LC simulation) is consistent with the literature value (2.30 nm^2^/ns^55^), and its relative contribution to the propagators is also compatible with the number of water molecules in the bulk region (Table S2). The second mode is significantly slower, by a factor of 5 from the bulk diffusion, very close to the 6-fold decreased diffusivity reported for buffer molecules within FUS condensates.^22^ Based on this agreement, and its contribution to the propagator, we expect *D*_2_ reflects slower water diffusion inside the protein-rich region (Table S2). The slowest mode (∼0.8% contribution) is difficult to pinpoint but is likely coming from a combination of factors, most importantly, water molecules directly interacting with protein atoms (Table S2). Similar to water diffusion, the dynamical behavior of ions reflects the presence of distinct populations. Most importantly, each mode’s contribution and its relative difference from bulk diffusion appear to depend on the protein sequence (Table S2).

The protein dynamics inside the condensed phase is closely connected to its liquid-like properties, needed for maintaining the biological function of the biomolecular condensate.^56^ To estimate the rate of relaxation of intramolecular protein degrees of freedom, we calculate 1.2 the time autocorrelation function of the radius of gyration (Fig. S5) yielding average correlation times of 192 and 122 ns for FUS LC and LAF-1 RGG respectively. We note that relaxation timescales for LAF-1 RGG are somewhat shorter compared with those for FUS LC, and that they are comparable to experimental estimates for isolated IDPs of similar length.^57,58^ This suggests that formation of the condensed phase has only a modest effect on intramolecular dynamics. The 2 *µ*s long MD simulations are at least 10 times longer than this relaxation timescale, which gives reasonable confidence in our ability to directly compute the diffusivity values of these two proteins (Fig. 3). The diffusion coefficient obtained for FUS LC is in excellent agreement with a previously determined value from FRAP and NMR diffusion experiments by Fawzi and co-workers^4,22^ (Fig. 3). Consistent with the faster chain relaxation time, the LAF-1 RGG diffusion coefficient is higher than that for the FUS LC. This may be explained because both the interchain and intrachain interactions governing frictional effects should have a similar dependence on the protein sequence.

**Figure 3.**
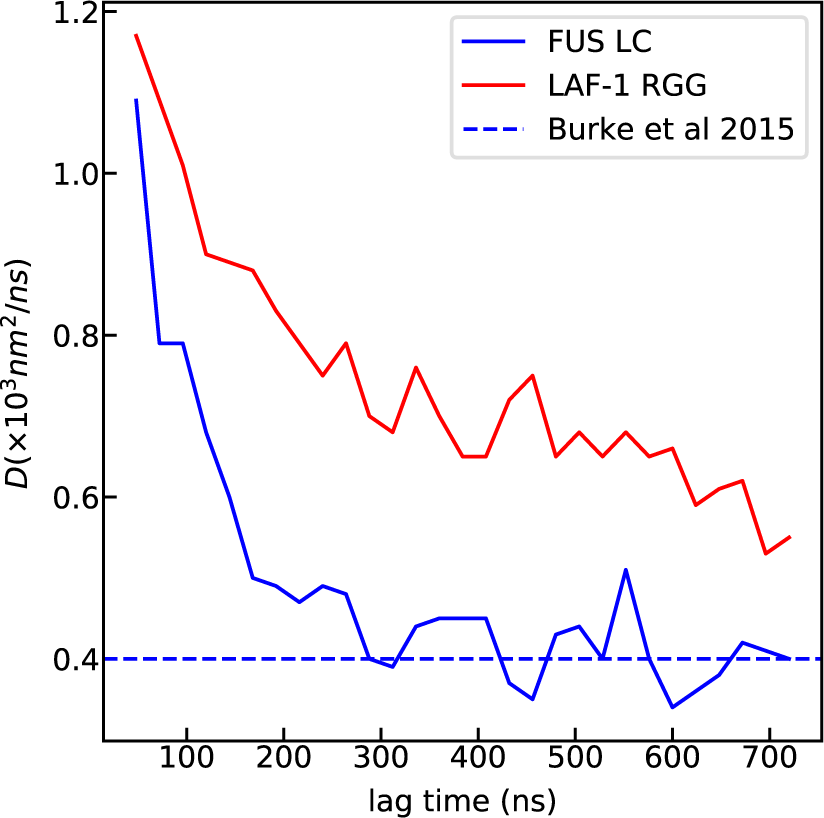
Protein self-diffusion coefficients along z-axis from all-atom slab simulations of FUS LC (blue) and LAF-1 RGG (red) are shown as a function of the lag time. Dashed horizontal line indicates experimentally determined diffusivity value for FUS LC.^4^

Because of the potential significance of secondary structure elements in mediating interactions in condensed phases,^44^ we examined the secondary structure populations of the proteins in the condensed phase using DSSP^59^ (Fig. S6). From this analysis, we find that the protein chains are largely disordered, with more than 50% of residues in a coil conformation, with local helices being the most common type of structured state (Fig. S6). This is consistent with experimental NMR studies showing a lack of structure within FUS condensates,^4,22,29^ and condensates of a protein similar to LAF-1 IDR, Ddx4.^52^

The central goal of this work was to elucidate the atomic-resolution interactions stabilizing a condensed proteinaceous phase which cannot be accessed through lower-resolution CG simulations. This information is essential to gain a fundamental mechanistic understanding of molecular driving forces and developing theory for the sequence determinants of protein assembly. Previous studies have highlighted the role of various interaction modes that may be responsible for driving the LLPS of different protein sequences, such as salt bridges,^52,60^ cation-*π* interactions,^61,62^ hydrophobic interactions,^22,38,63^ sp^2^/π interactions between several residue pairs including the protein backbone,^21,22^ and hydrogen bonding interactions.^22,38,64^ There is still a limited understanding of the relative importance of these different interaction modes in the context of a particular type of amino acid pair, or a protein sequence. We attempt to provide answers to some of these questions here.

To characterize the regions of each sequence most involved in molecular interactions, we start by computing the number of intermolecular van der Waals (vdW) contacts formed as a function of protein residue number (Figs. 4a and b) per frame, averaged over the entire trajectory. We find that contacts are relatively evenly distributed throughout the FUS LC sequence (Fig. 4a), which is consistent with NMR data.^22^ One can observe intermittent peaks in the one-dimensional contact map data arising from the Tyr residues distributed throughout the FUS LC sequence (Fig. 4A, black dashed lines in the bottom panel). For LAF-1 RGG, the contacts are still distributed throughout the chain with a notable contact-prone region between residues 20-28 (Fig. 4b), which was identified previously from our CG model simulation and tested experimentally to be critical for promoting LLPS.^35^

**Figure 4.**
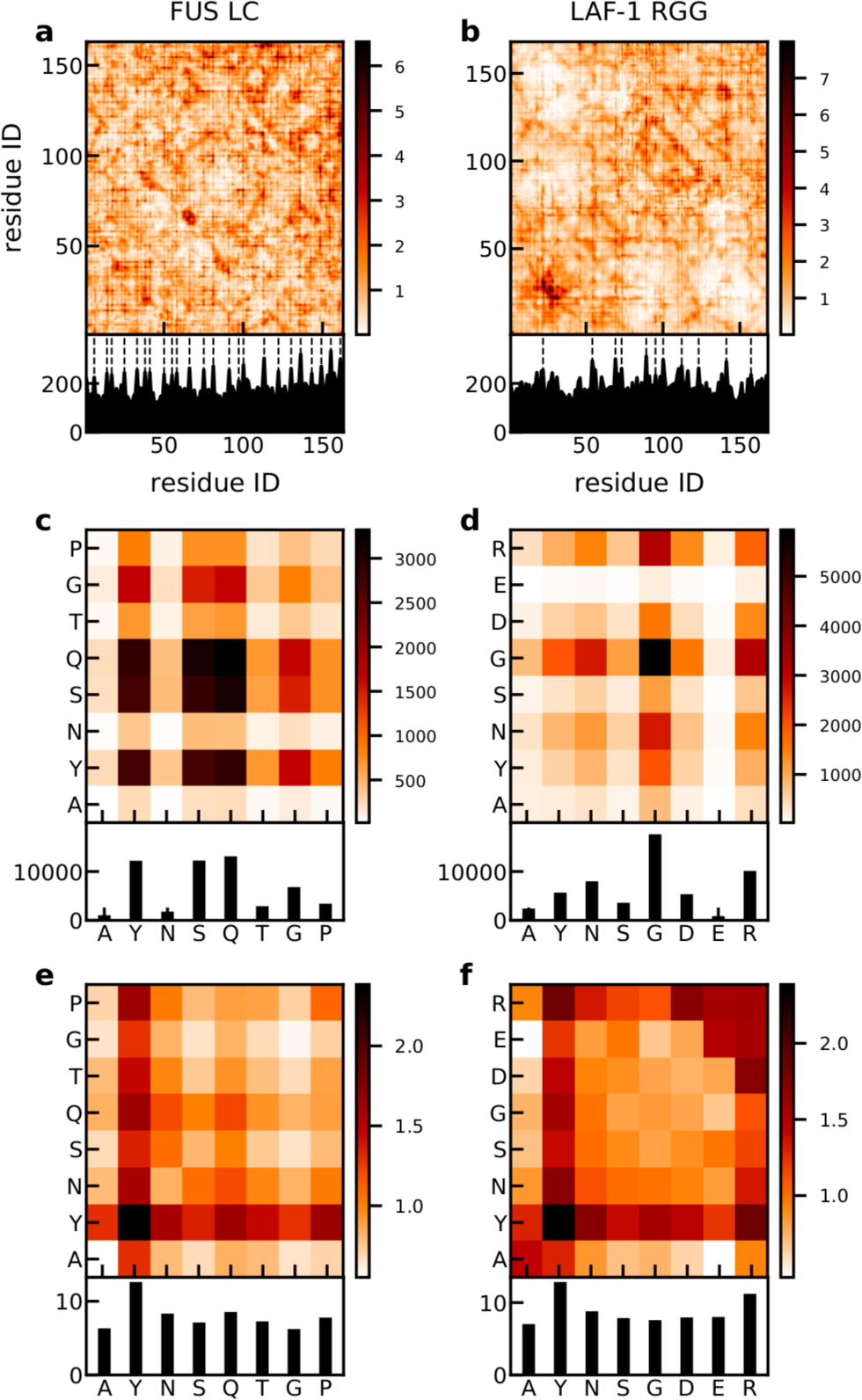
Intermolecular contacts within the condensed phase of FUS LC (left) and LAF-1 RGG (right), as a function of residue index (a and b) and amino acid types (c and d). The intermolecular contacts normalized by the relative abundance of each amino acid in the sequence are shown in e and f. Black dashed lines in a and b show the position of Tyr residues.

To obtain a better understanding of how different amino acid types contribute to the formation of intermolecular contacts between protein chains, we combine the contact data for different pairs of the same kind. From this data in Figs. 4c and d, one can identify important residue pairs as well as residue types that are primarily responsible for interchain interactions and for stabilizing the condensed protein-rich phase. For both FUS LC and LAF-1 RGG, Tyr interactions with itself and other residues are highly abundant and likely essential drivers of LLPS.^21,22,36,60,62,64,65^ Importantly, polar residues (Gln and Ser in the case of FUS LC and Asn in LAF-1 RGG) also participate in significant contacts, consistent with a recent mutagenesis study highlighting their role in LLPS.^22^ Also, Gly residues appear to be forming contacts with many other residue types in both proteins; this is highly visible in the LAF-1 RGG data for interactions with Arg, Gly, Tyr, and Asn. Lastly, LAF-1 RGG contact formation is enhanced by interactions between oppositely-charged residue pairs such as Arg-Asp pairs (Fig. 4d). To obtain the intrinsic propensity for each amino acid to form a contact, we also normalize the contacts by the relative abundance of each amino acid in the sequence (Figs. 4e and f). The overall values are largely consistent between the FUS LC and LAF-1 RGG simulations, and supports the critical role of Tyr and Arg due to their intrinsic preference to form contacts while the Gly-involved contacts are present due to its abundance in both the sequences. Even though Gln and Ser contribute as many total contacts as Tyr, each individual Tyr contributes more. This is because there are more Ser and Gln than Tyr in FUS LC sequence. Additionally, Tyr is bigger than Ser and Gly, which may allow it to make more simultaneous contacts. We have also calculated the intramolecular interactions in the same way and obtained quantitative agreement with the intermolecular interactions (Figs. S7 and S8), which supports our recent finding connecting the self-interaction properties of the single chain with LLPS behaviors.^31^ Data for residues that are an insignificant fraction of the protein composition (appearing ≤2 times) have been excluded from the plot due to higher uncertainty associated with their contacts.

To dive deeper into the atomic interactions responsible for the observed role of the amino acids identified above, we determine the interaction modes present when two residues form a vdW contact. Based on the previous literature,^21,22,62^ the most important modes are sp^2^/*π*, hydrogen bonding, cation-*π*, and salt bridge. Here, we separate these interactions into contacts between backbone atoms (bb-bb), sidechain atoms (sc-sc), or backbone and sidechain (bb-sc) atoms (see supporting methods for definition of these interaction modes). The amino acid pairs are sorted by the number of vdW contacts formed (Fig. S9) and the top 20 amino acid pair types for each group and each protein are shown in Fig. 5 with the full version in Fig. S10. The interaction modes from intramolecular interactions (Figs. S11 and S12) are highly similar to those from intermolecular interactions, so we only discuss the intermolecular version here.

**Figure 5.**
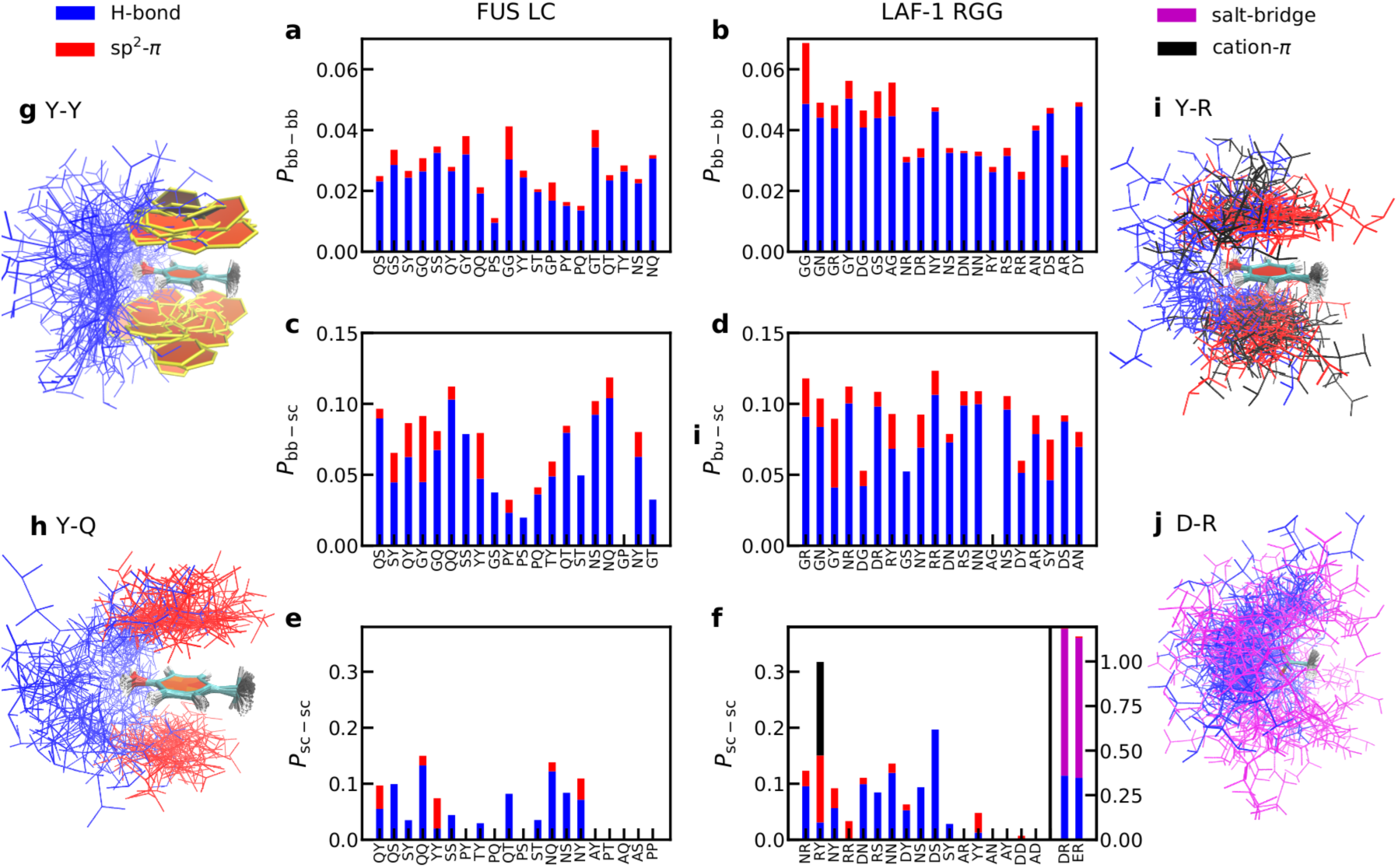
Interaction modes contributing to the intermolecular contacts separated into contributions between backbone atoms (bb-bb, a and b), between backbone and sidechain atoms (bb-sc, c and d) and between sidechain atoms (sc-sc, e and f). The amino acid pairs are sorted by the number of contacts formed between these pairs (Fig. S9) in each group. Some configurations of representative interacting amino acids are shown in g for FUS LC Tyr:Tyr, h for FUS LC Tyr:Gln, i for LAF-1 RGG Tyr:Arg and j for LAF-1 RGG Asp:Arg. The configurations are aligned to Tyr in g, h and i and to Asp in j. The color code is the same as the interaction mode shown in the legend. Rings are shown in paperchain representation.^66^

For both FUS LC and LAF-1 RGG, we first note that most of the vdW contacts are non-specific and thus only a small fraction can be classified into any of the aforementioned specific interaction modes (Figs. 5a–f). This emphasizes the importance of non-specific interaction modes, including the hydrophobic interactions, in promoting LLPS. Within the context of interaction modes in FUS LC, all pairs except for those involving Tyr are primarily stabilized by hydrogen bonds(Figs. 5a,c and e). Surprisingly, interactions involving sp^2^/*π* groups are a relatively small fraction of the contacts, except for residue pairs involving Tyr, with contributions from the sp^2^/*π* mode higher than from hydrogen bonds. The configurations of representative amino acid interactions are also shown in Figs. 5g and h. For both Tyr and Gln interactions, sp^2^/*π* interaction modes tend to form on top or bottom of the sidechain whereas hydrogen bonds are around the side. This suggests for aromatic amino acids like Tyr, hydrogen bonds might not directly compete with forming sp^2^/*π* interactions and can still be a major contribution to stabilizing the condensates.

For the LAF-1 RGG, we include two additional interaction modes, salt bridges and cation-*π* interactions, involving charged residues (Figs. 5b,d and f). We find that charged amino acids contribute heavily to LAF-1 sidechain interactions, with hydrogen bond and sp2/*π* interactions from aromatic amino acids playing secondary roles (Fig. 5f). Previously, we have shown that certain pairings of residues can form contacts using different interaction modes, either switching between them, or forming multiple contacts cooperatively, ^35^ particularly cation-*π* and sp^2^/*π* interactions between Arg and Tyr. Here we also find hydrogen bonds and salt bridges between cationic-aromatic pairings and oppositely-charged residues are among the strongest interactions occurring within the LAF-1 condensate (Fig. 5f), and different interaction modes can occur at the same time, e.g. cation-*π* and sp^2^/*π* interactions between Arg and Tyr (Fig. 5i), and salt bridge and hydrogen bonds between Arg and Asp (Fig. 5j). This is also the reason the total probability of interaction modes might exceed 1 for interactions involving charged amino acids (Fig. 5f).

## Conclusion

In this work, we present a general methodology for initializing, conducting, and analyzing all-atom explicit-solvent simulations of biomolecular condensates in coexistence with a surrounding aqueous phase. We have optimized the procedure for systems with components of similar size to FUS LC and LAF-1 RGG so that similar simulations should be accessible using even general-purpose computing hardware (Table S1). We have leveraged our earlier work with CG simulations of IDP phase coexistence,^5,22,24,29,31,35,40,41,67^ and atomistic studies of inter-protein interactions^5,22,24,35^ to obtain important mechanistic details of the underlying molecular interactions of condensates.

We find that the proteins are remarkably dynamic in the condensed phase, having intramolecular correlation times very comparable to those typical of isolated intrinsically disordered proteins. This flexibility is key to the liquid-like properties of the protein-rich phase. While the dense phase is highly viscous, we are also able to measure the protein diffusivity, finding excellent agreement with experimental results where available. Similarly, we show that water and ions are able to rapidly diffuse between phases, with diffusion coefficients within the dense phase reduced.

For both tested proteins, the equilibrium distribution of sodium and chloride ions within the condensed phase is essentially determined by the charge distribution and water content inside the phase-separated proteins. This implies that there is no strong preferential interaction of these ions with protein residues in these systems under the conditions we study. We note, however, that ions exhibiting stronger Hofmeister effects, or higher salt concentrations,^68–70^ may alter this result, and would be interesting to consider in future work.

Finally, we find many types of residue-residue interactions are responsible for stabilizing the condensed phase, and contacts involving Gly are particularly abundant due to its frequency in the sequence. After normalizing for residue frequency, however, it appears that each Tyr contributes more interactions per residue than any other residue type, explaining its apparent importance in mutagenic approaches. For LAF-1 RGG, in addition to Tyr interactions, we observe that both cation-*π* interactions (particularly involving Arg) and salt bridges contribute to the condensate’s stability. The approach outlined here can be used to explore the generality of these findings in the context of other protein sequences.

## Acknowledgement

We acknowledge useful discussions with Dr. Anastasia Murthy. This work was supported in part by the National Institutes of Health grants R01GM118530 (N.L.F.), R01NS116176 (N.L.F. and J.M.), and R01GM120537 (J.M.), National Science Foundation grants DMR-2004796 (J.M.) and MCB-2015030 (W.Z.). R.B. was supported by the Intramural Research Program of the National Institute of Diabetes and Digestive and Kidney Diseases of the National Institutes of Health and Y.C.K by the Office of Naval Research via the U.S. Naval Research Laboratory base program. This research used computational resources of Anton 2, XSEDE (supported by the NSF project no. TG-MCB120014), and the NIH HPC Biowulf cluster (http://hpc.nih.gov).

## Supporting Information

### Supporting Methods

#### Setup of all-atom slab simulations

The challenge in modeling the condensed phase of an IDP at atomic resolution is to equilibrate the density of the condensed phase. Even using the coarse-grained (CG) model, it takes about 1 *µ*s simulation time to achieve the task.^S1^ Here by using all-atom model, this is expected to be much slower considering the explicit solvent and a much greater number of degrees of freedom involved in the simulation. We therefore explore an alternative approach by initializing the all-atom simulation from a reasonable well-equilibrated CG conformation.

Using FUS LC as an example, we first generated an initial configuration of the FUS chains with our recently developed CG model–HPS model.^S1^ We followed the same protocol used in the CG modeling by setting up a slab geometry, in which one box dimension is elongated, allowing for a semi-infinite condensed phase in two dimensions, and two flat interfaces along the elongated dimension (see Fig. 1A). This geometry has been widely used,^S2,S3^ and demonstrated to be comparable with other accepted methods of sampling multiple phases, such as grand canonical Monte Carlo^S4^ while reducing the effects of a finite-sized spherical droplet.^S5^ The other advantage of the slab geometry in the case of explicit solvent is that it greatly reduces the box volume that is occupied largely by aqueous phase, compared to a brute-force droplet simulation. To reduce the overall box size, we have conducted CG simulations of a slab of FUS chains containing a different number of IDP chains, and at different box cross sections in order to obtain a system that has a sufficient fraction of the chains in the condensed phase region, rather than interfacial region (Fig. S1). The number of chains (40 here) included in the simulations have been carefully selected so that it is large enough to still reproduce exactly the density profile of our original CG simulations with 100 chains at the same temperature, whereas at the same time to be as small as possible to reduce the system size so that the simulation is still feasible.

With a reasonable slab configuration in CG representation as the starting point, we further prepared the all-atom configuration following the steps shown in the main text and Fig. 1. Finally, the system is solvated with explicit water molecules and ions, and further equilibrated using standard molecular dynamics engines (Fig. 1D). The equilibration is conducted using GROMACS 2018.^S6^ Langevin dynamics was performed with a time step of 2 fs and a friction coefficient of 1 ps^−1^. Berendsen pressure coupling was used for 10 ns followed by Parrinello-Rahman pressure coupling^S7^ for 200 ns simulations. Lennard-Jones (LJ) pair interactions were cut off at 0.9 nm. Electrostatic energies were computed using particle-mesh Ewald^S8^ with a grid spacing of 0.12 nm and a real-space cutoff of 0.9 nm. The protein force field was Amber ff03ws;^S9^ the water model was TIP4P/2005^S10^ and the ion parameters were from Luo et al.^S11^ The final system we prepared for FUS contains 40 chains of the 163-residue protein, solvated in 98912 water molecules,^S10^ 280 Na^+^ ions and 200 Cl^−^ ions to neutralize the system net charge, and represent an ionic strength of about 100 mM, with a total of 485408 atoms (see Table S1).

Further 150 ns equilibration simulations using an NPT ensemble were conducted on Anton 2–a specialized supercomputer^S12^ together with two production simulations, one for FUS LC and one for LAF-1 RGG for a total of 2 *µ*s each using the same aforementioned force fields. We need to note that there have been a series of force fields developed for IDPs and can be used for similar simulations.^S9,S13–S15^ The Amber ff03ws force fields have been demonstrated to capture the configurations and dynamics of disordered proteins.^S16,S17^ It should be noted that performance on traditional resources, while one or two order of magnitude slower than on Anton 2, should be sufficiently fast to generate microseconds worth of data within a reasonable time frame. For instance we obtained the initial equilibration with a benchmark of about 45 ns/day with 896 Xeon E5-2680 CPUs. This makes such a 2 *µ*s simulation feasible even using traditional computational resources. Please see Table S1 for simulation details and benchmarks.

### Sequences of the proteins used in this work

N-terminal low complexity domain of the FUS protein (FUS LC)

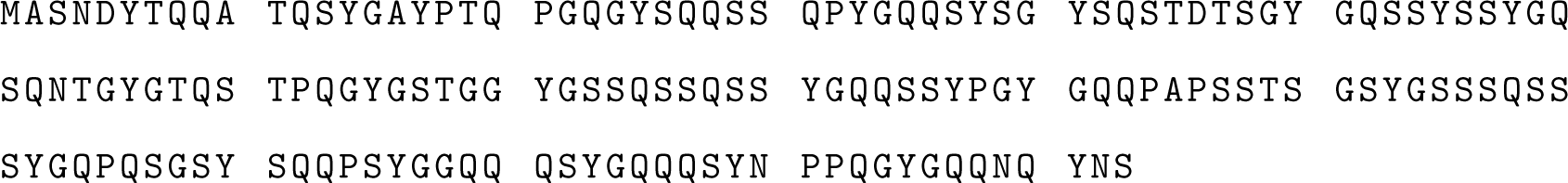

N-terminal disordered domain of the LAF-1 protein (LAF-1 RGG)

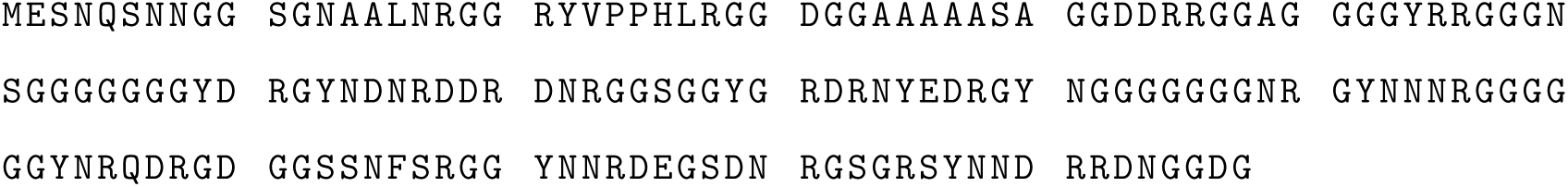

### Calculation of diffusion coefficients

To estimate the diffusion coefficients in the simulation, we first compute the probability distributions (propagators) *P*(*ξ*(*t* + *t*_0_) − *ξ*(*t*_0_)) for molecular displacement in each direction *ξ* = *x, y, z* as a function of the lag time *t* between observations to accurately describe the dynamics of protein, water, and ions (i.e. Na^+^ and Cl^−^). We then fit the probability distribution function at a specific lag time to the 1D diffusion equation with multiple diffusivity values as

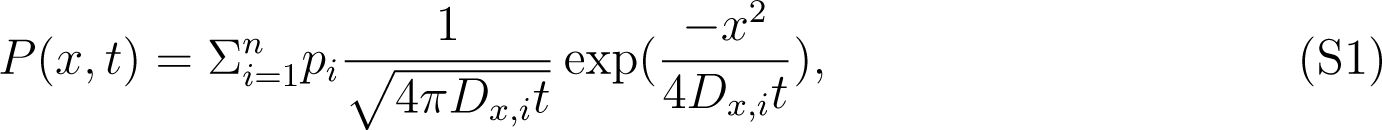

and

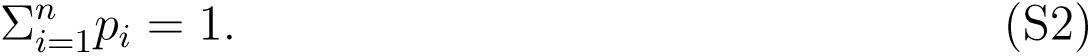

We increase the number (*n*) of diffusivity (*D*) values needed for fitting the probability distribution until the fitting is reasonably good as shown in Fig. S4. We find that for the protein, one *D* value (*n* = 1) is sufficient whereas for the water and ions, we need three *D* values (*n* = 3) for some of the lag time. We then check *D* as a function of lag time until the *D* values are plateaued at a range of the lag time. The average values of *D* at that window are reported (Table S2). The lag time window for calculating the protein diffusion coefficients are from 408 to 720 ns and that for water and ions are from 3.6 to 6 ns.

### Extracting molecular interactions from all-atom simulations

#### Contact formation

A contact between two groups (i.e. backbone or sidechain of amino acids) are considered if at least one atom from each of the two groups are within 6 Å distance.

#### Hydrogen bond

MDAnalysis is used in calculating the hydrogen bond formation^S18^ with a distance cutoff of 3.0 Å between the donor and acceptor and an angle cutoff of 120^◦^ between donor, hydrogen and acceptor.

#### sp^2^/π interaction

We calculated the sp^2^/*π* interactions based on the definition of a recent literature^S19^ with small modification of the algorithm for efficiency. First we filtered all the pairs of sp^2^/*π* groups by using a cutoff of 8 Å on the distance between the center of mass of the two groups. Second we calculated cosine angles between the normal vectors of the two plains and only kept the groups with absolute values of cosine angles larger than 0.8. Third both the plains defined by each group were raised by 1.5 Å and the distance between the center of mass of the two new plains were calculated. The pairs with the center of mass distances less than 4 Å were selected as forming the sp^2^/*π* interactions.

#### Cation-π interaction

Similar to the sp2/*π* interactions, Cation-*π* interactions are also defined by using both a distance and an angle criterion. The distance between the charged nitrogen in the cationic side chain and the center of mass of the *π* group is first subjected to a cutoff of 6 Å. The absolute cosine angles between the normal vector of the *π* plain and the vector linking the charged nitrogen and the center of mass of the *π* group is further subjected to a cutoff of 0.8.

#### Salt bridge

We used a distance cutoff of 6 Å on the smallest distance between all charged nitrogen and oxygen atoms for every pair of charged amino acids to determine the formation of the salt bridge in our simulations.

---

**Figure S1.**
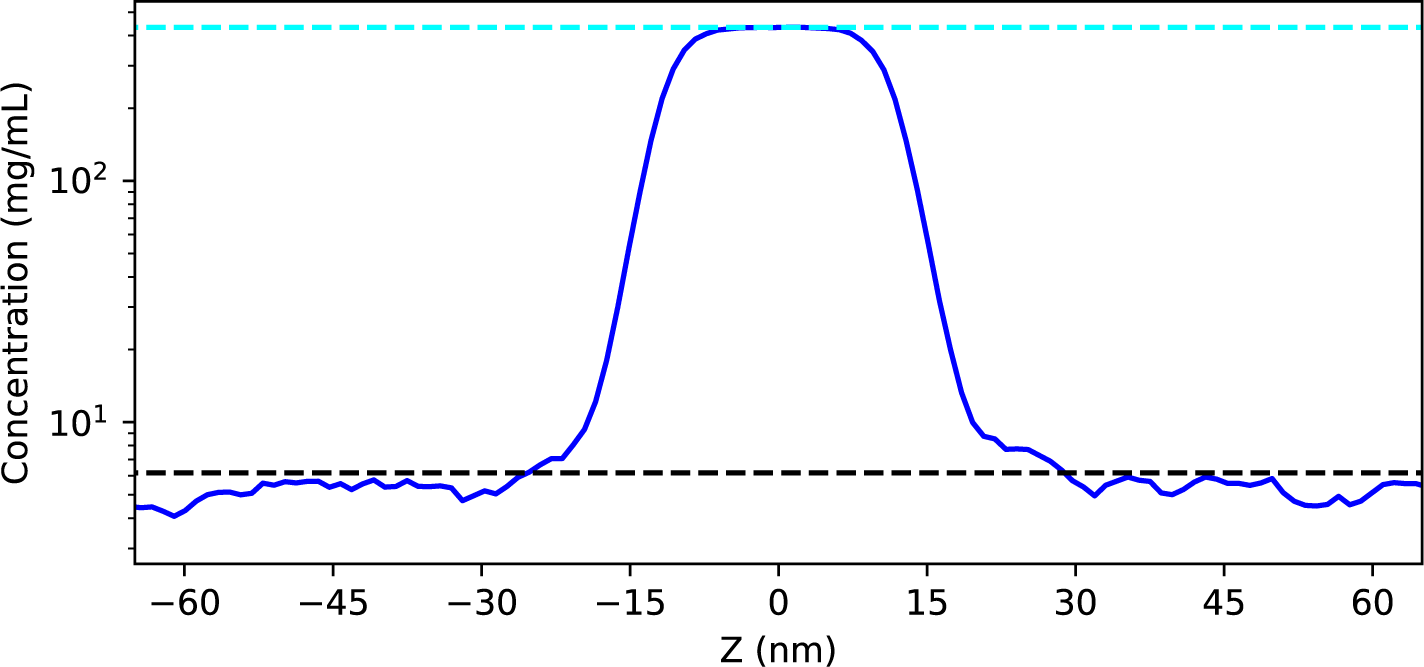
Test of CG simulations of FUS LC with 40 chains in slab geometry using a 10 nm box cross section at 340K compared to reference densities from a 100-chain slab simulation with a 15 nm box cross section as used in references.^S1,S5^ The black dashed line indicates the low density phase of the reference and cyan dashed line indicates the high density phase of the reference.

**Figure S2.**
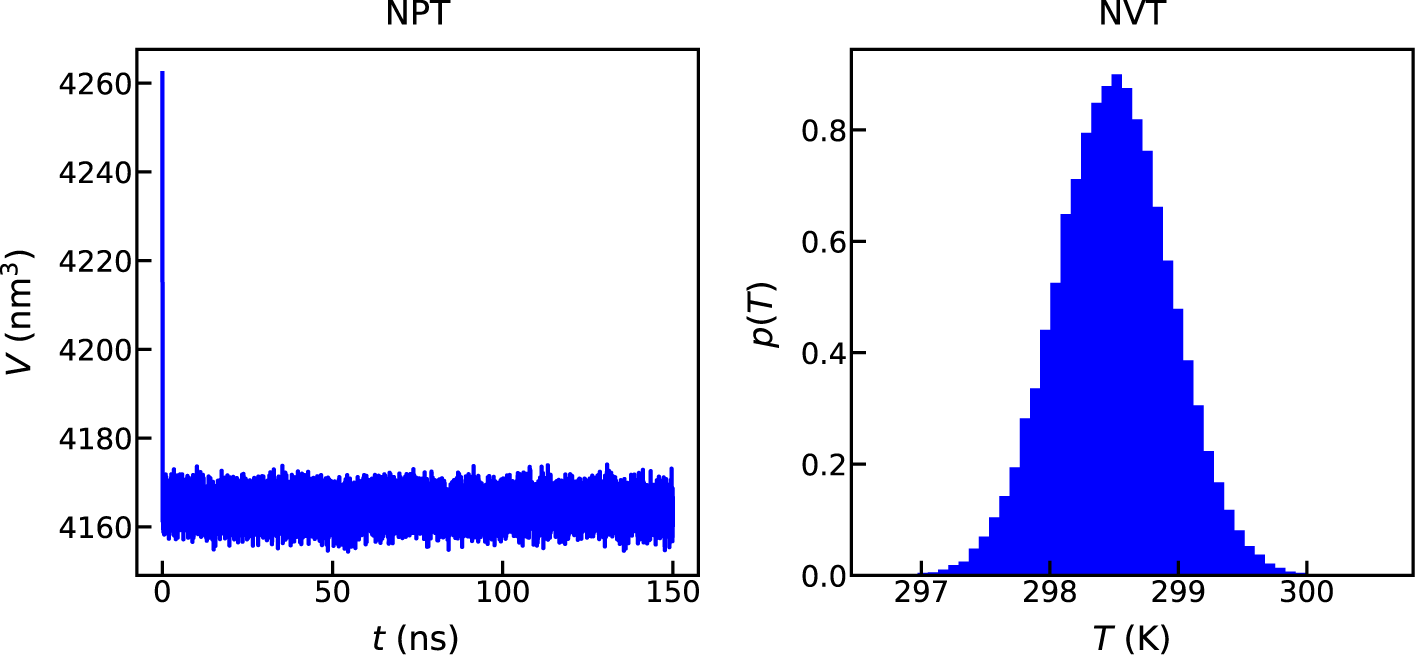
The volume fluctuation of NPT equilibration and temperature distribution of productive NVT simulation for LAF-1 RGG using Anton 2.

**Figure S3.**
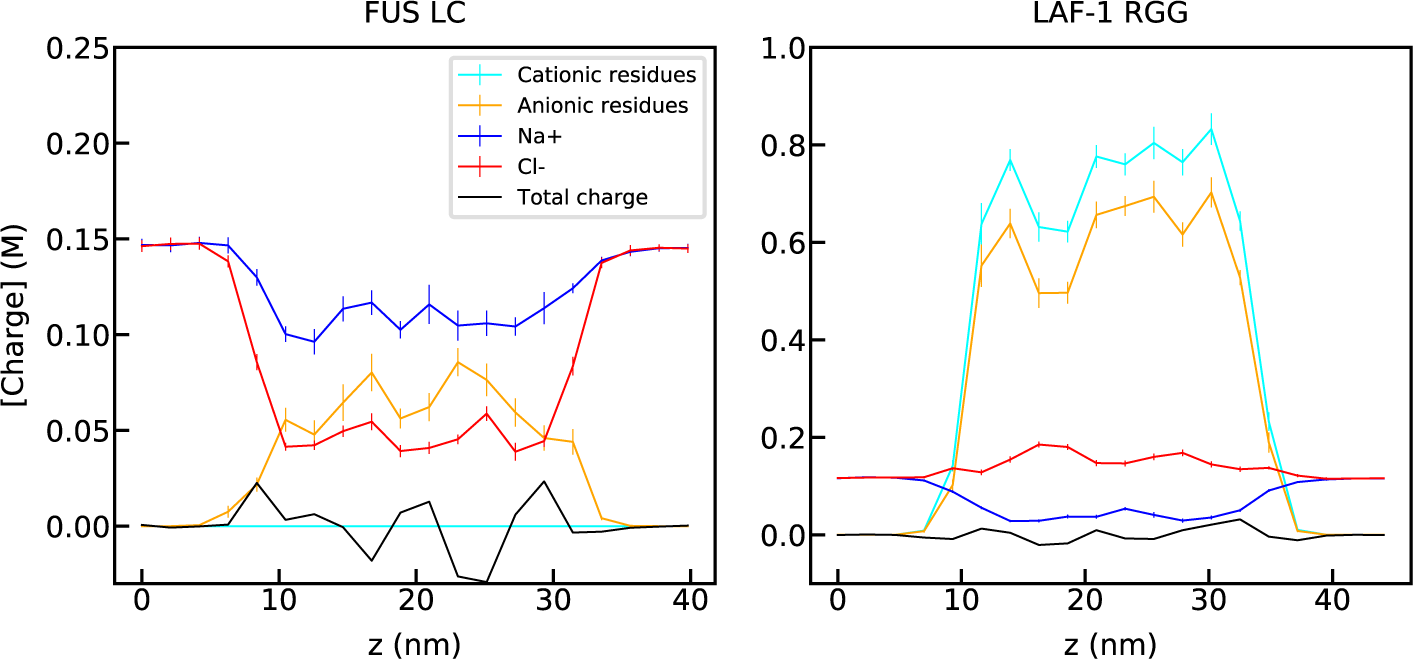
Density profiles of charges from all-atom slab simulations of FUS LC (left) and LAF-1 RGG (right). Components are shown in the legend.

**Figure S4.**
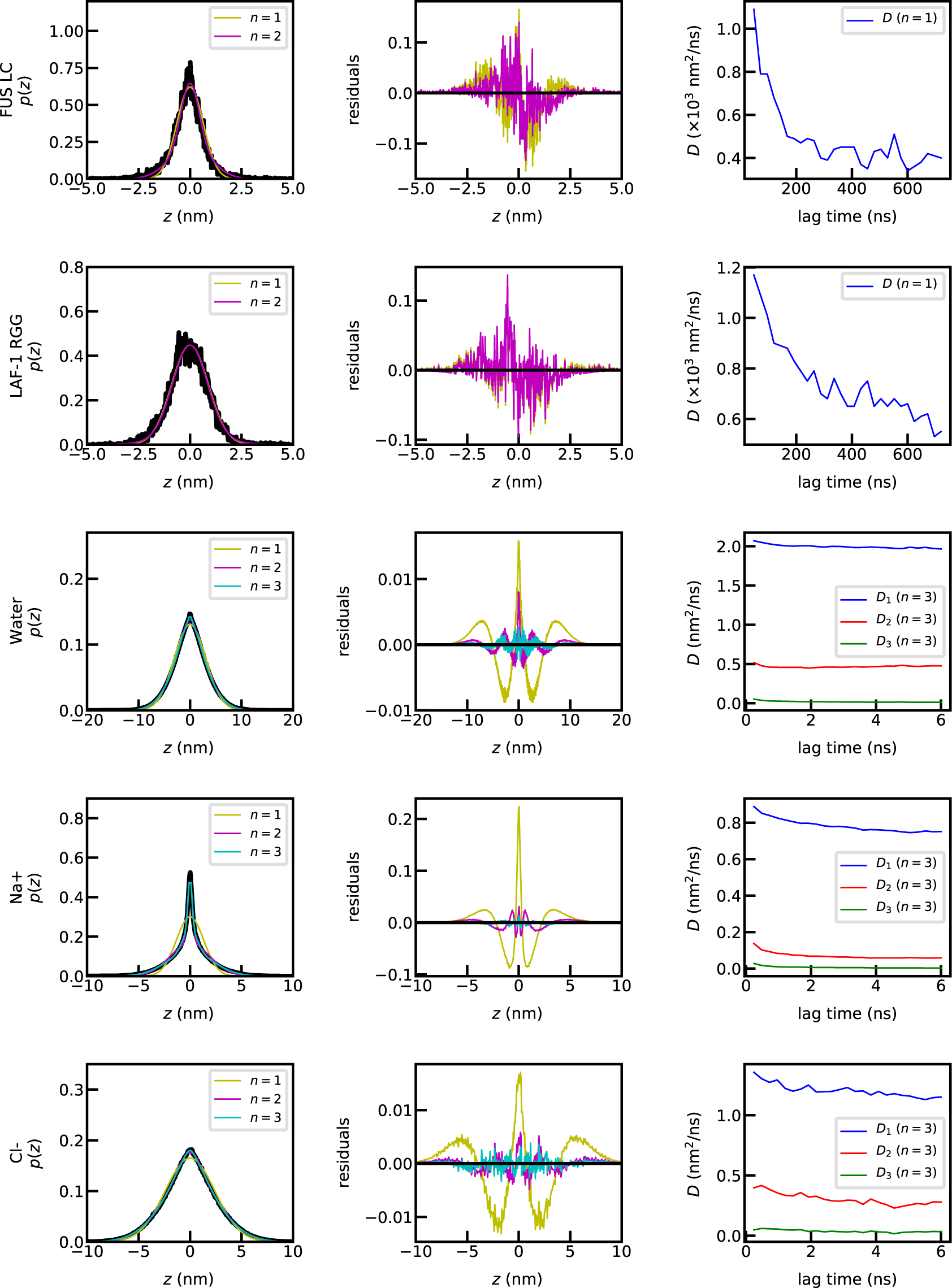
Representative distribution of mean square displacement at lag time of 600 ns for proteins and 3.6 ns for water and ions. The black lines show the results from simulations and the color lines show different strategies of fitting.

**Figure S5.**
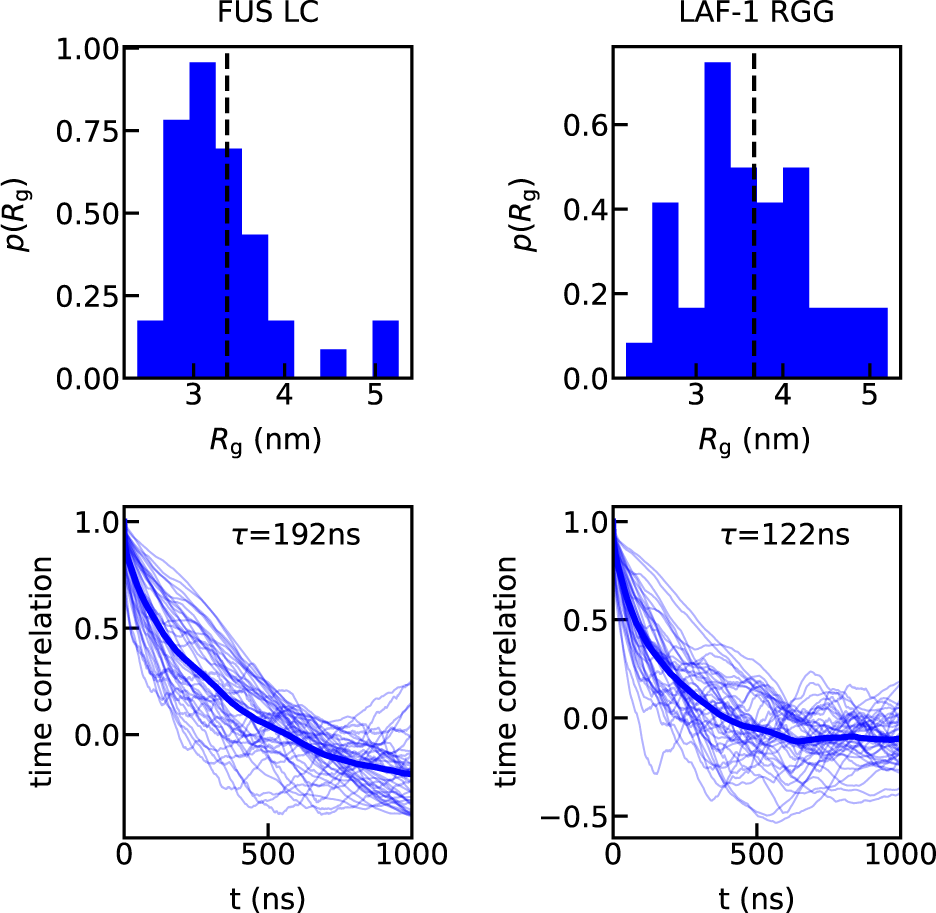
Radii of gyration (*R*_*g*_) of FUS LC (left) and LAF-1 RGG (right). Top: The distribution of the average *R*_*g*_ of each of the 40 chains in the simulation. Bottom: Time correlation of *R*_*g*_ for each of the 40 chains (thin lines) and the average over the 40 chains (thick line). The relaxation time obtained by fitting the average time correlation to a single exponential function is shown in the legend.

**Figure S6.**
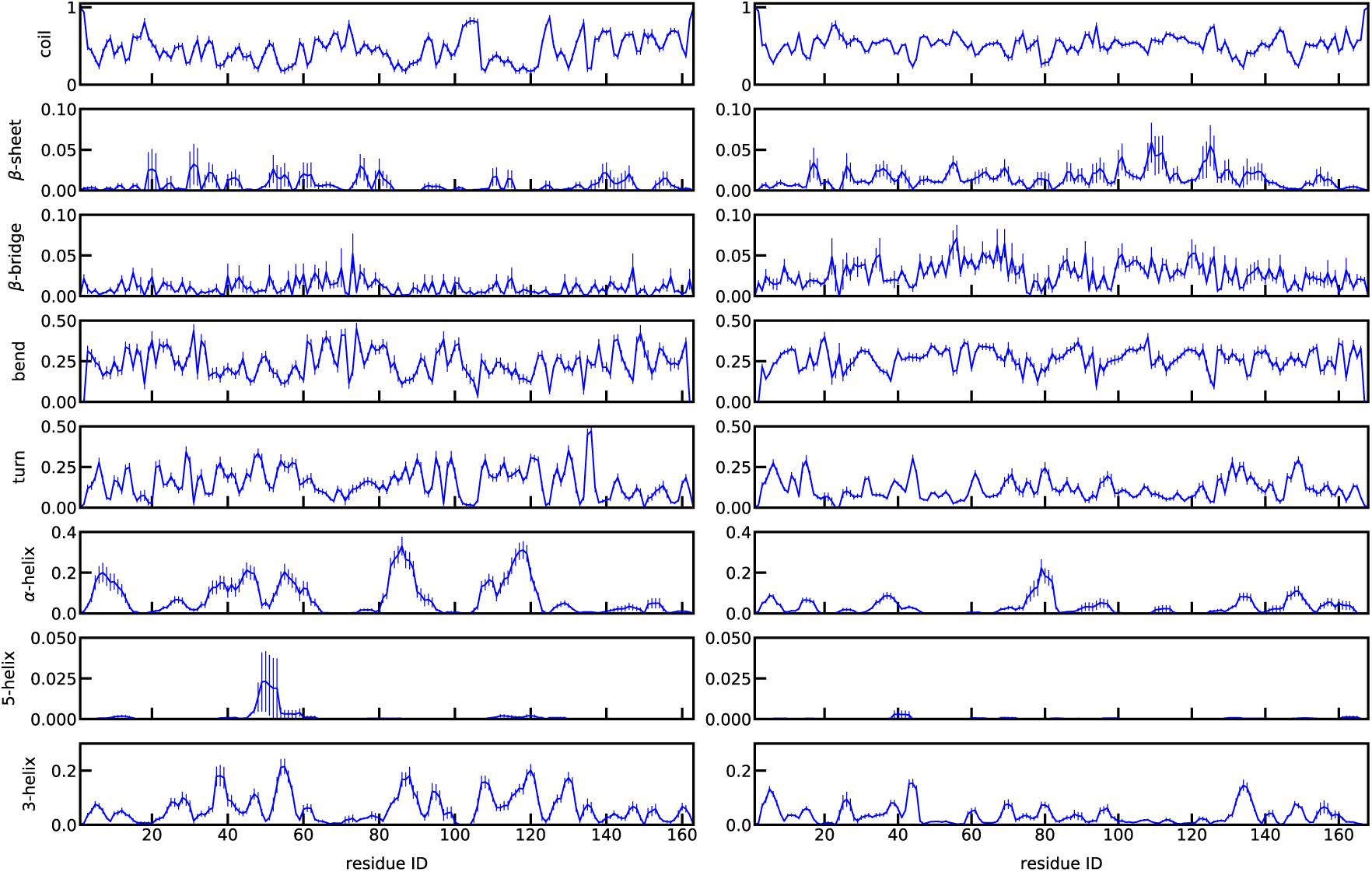
Secondary structures of FUS LC (left) and LAF-1 RGG (right) using DSSP.^S20^

**Figure S7.**
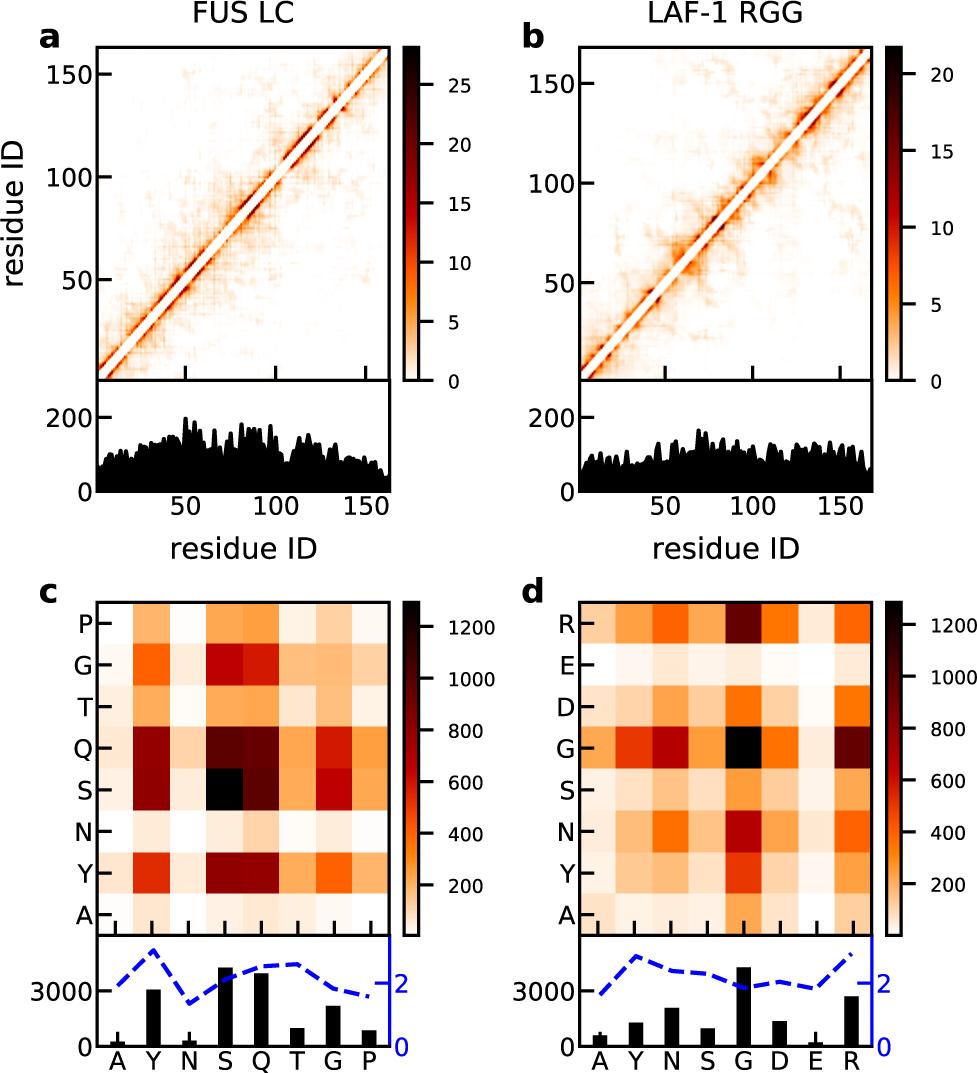
Intramolecular contacts within the condensed phase of FUS LC (left) and LAF-1 RGG (right), as a function of residue index (top) and amino acid types (bottom). Blue lines show the intramolecular contacts normalized by the relative abundance of each amino acid in the sequence in the sequence.

**Figure S8.**
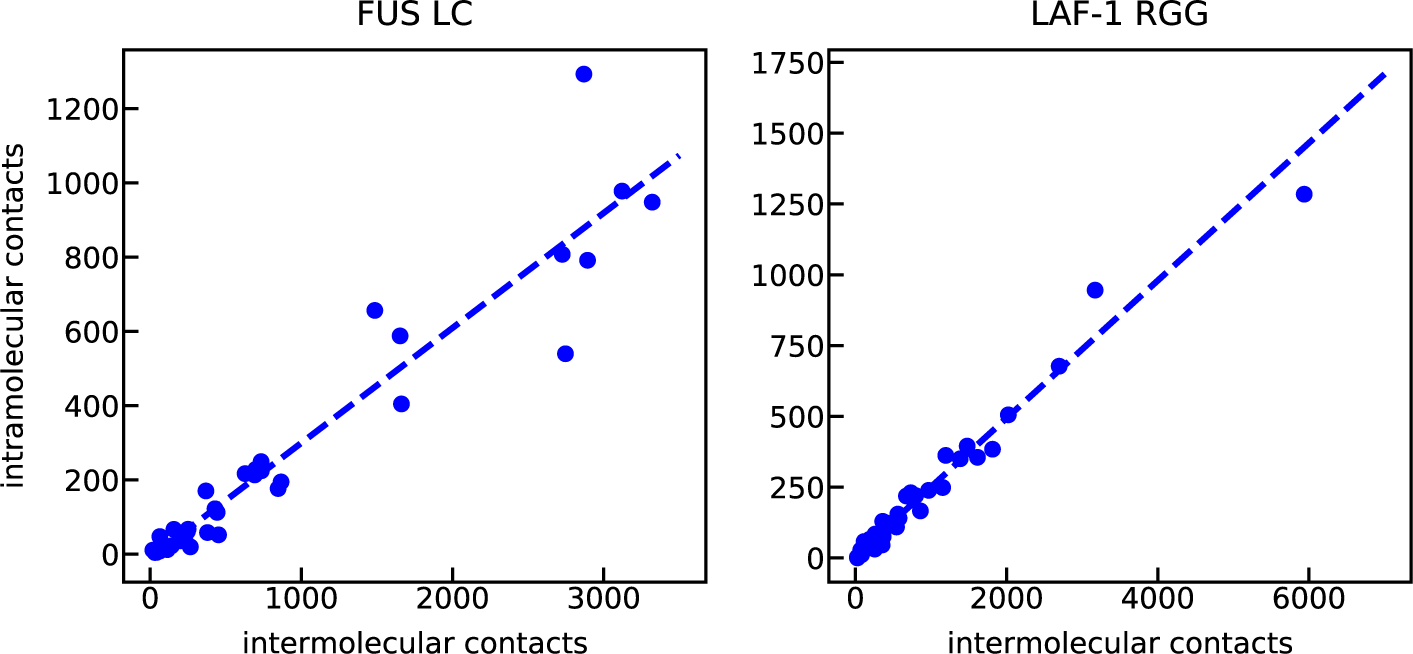
Correlation between intra- and inter-molecular contacts for each pair of amino acids within the condensed phase of FUS LC (left) and LAF-1 RGG (right).

**Figure S9.**
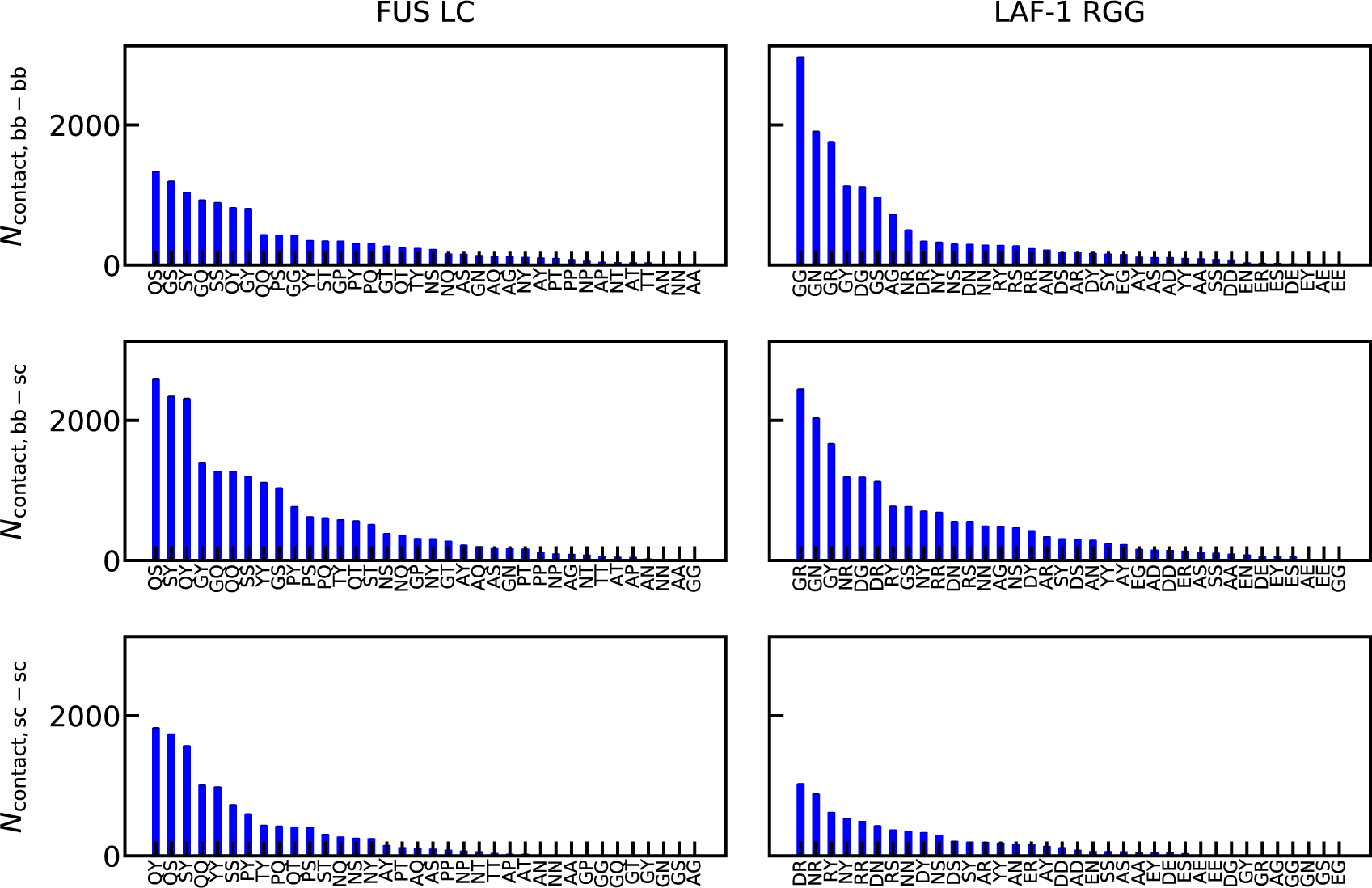
Number of intermolecular contacts for specific pairs of amino acids between backbone and backbone atoms (bb-bb), backbone and sidechain atoms (bb-sc), and sidechain and sidechain atoms (sc-sc).

**Figure S10.**
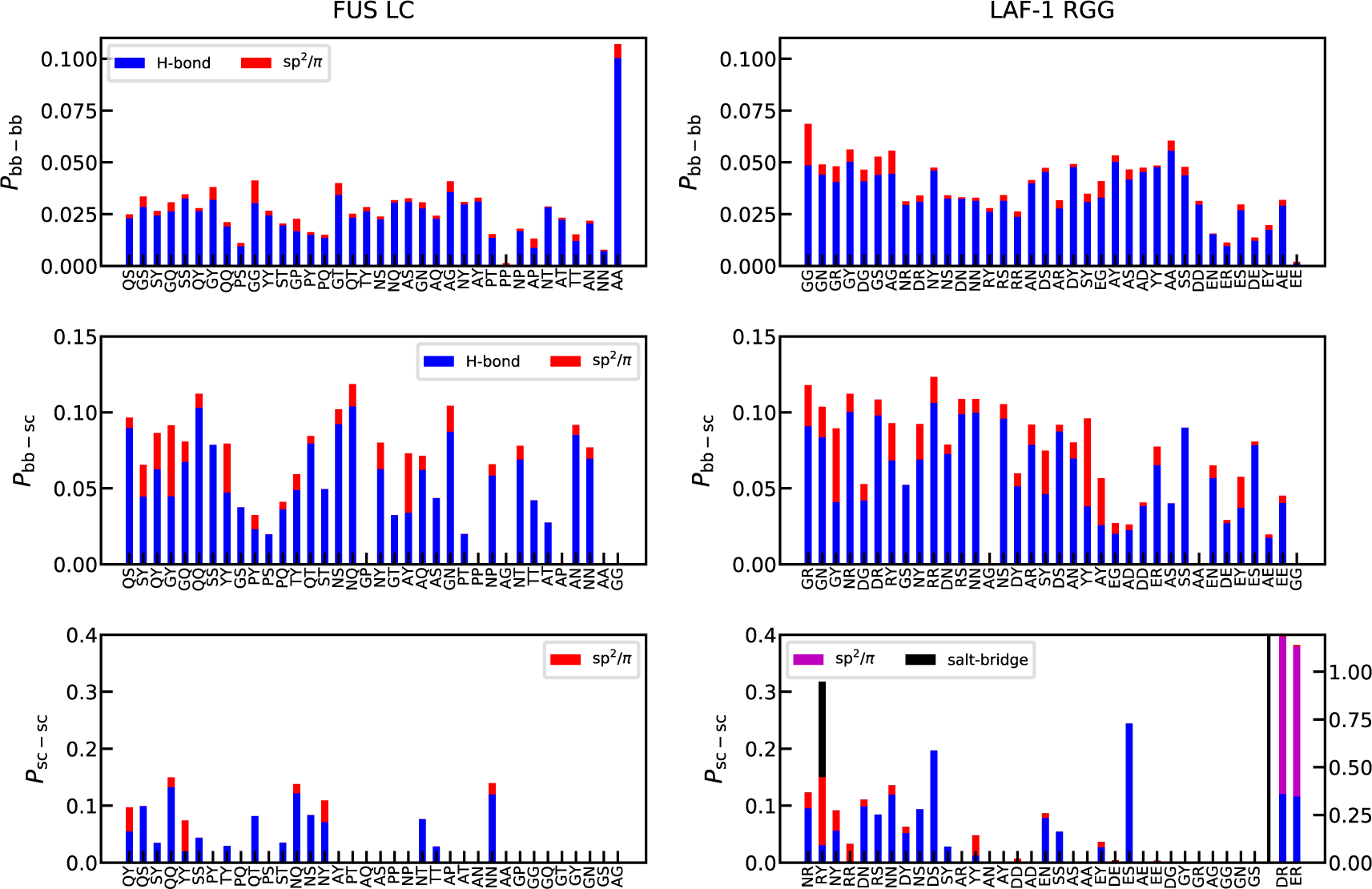
Interaction modes contributing to the intermolecular contacts.

**Figure S11.**
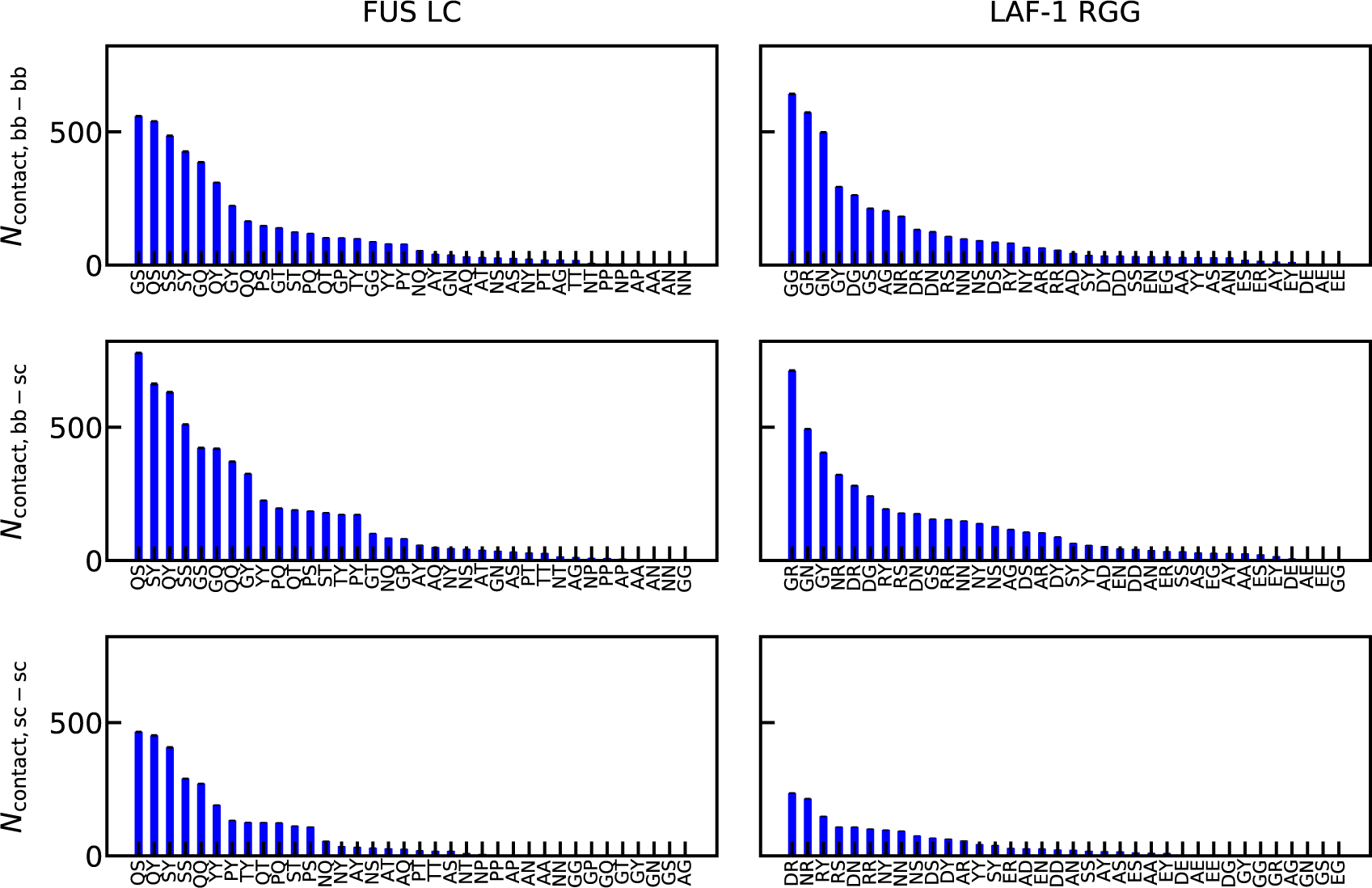
Number of intramolecular contacts for specific pairs of amino acids between backbone and backbone atoms (bb-bb), backbone and sidechain atoms (bb-sc), and sidechain and sidechain atoms (sc-sc).

**Figure S12.**
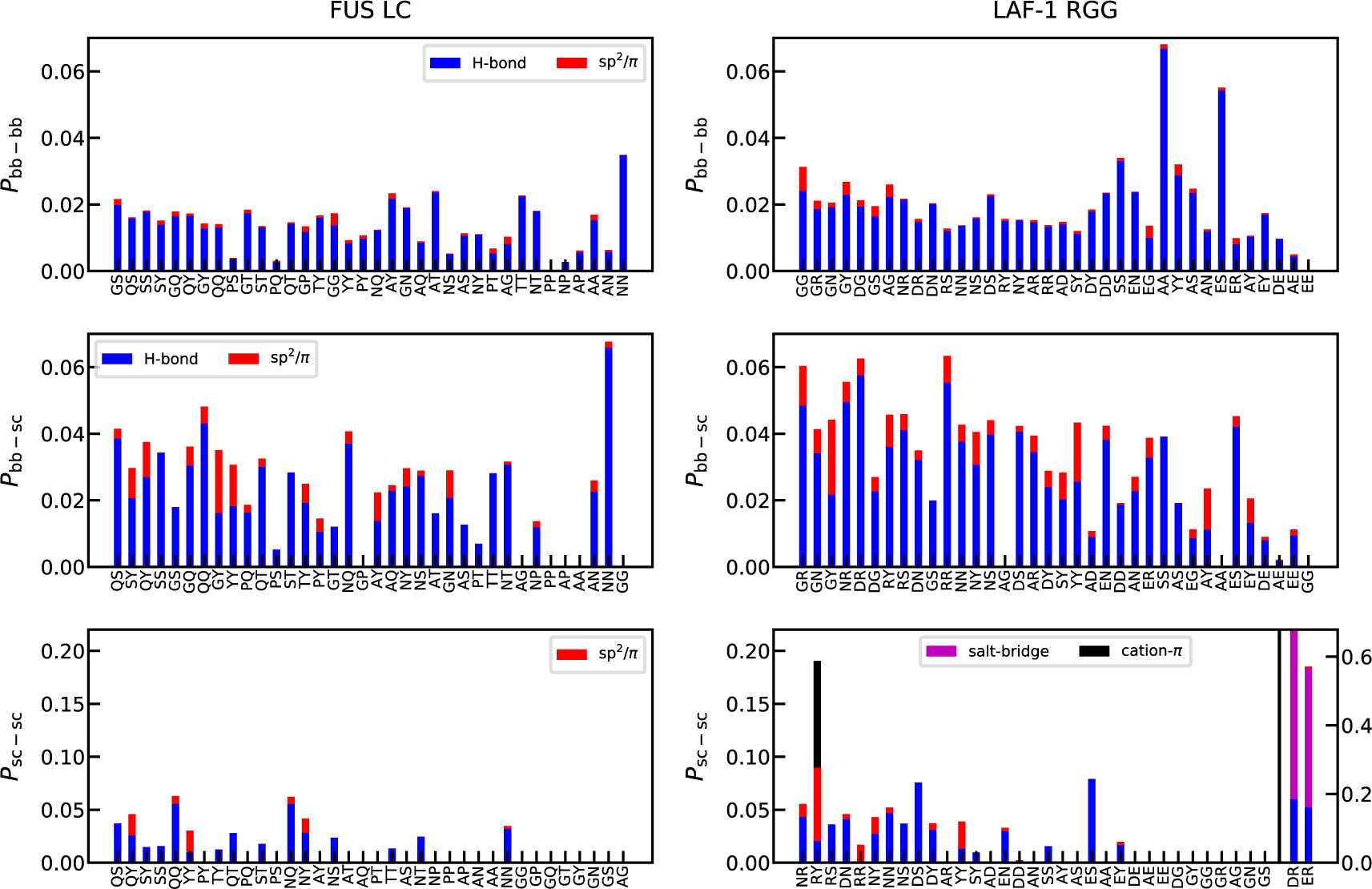
Interaction modes contributing to the intramolecular contacts.

**Table S1.** System sizes of all-atom simulations. *The benchmark of local simulations is estimated by using Gromacs 2018.3^S21^ and 896 CPUs of Xeon E5-2680.

| Name | FUS LC | LAF-1 RGG |
| --- | --- | --- |
| number of chains | 40 | 40 |
| number of amino acids per chain | 163 | 168 |
| number of water molecules | 98912 | 114727 |
| number of Na+ | 280 | 188 |
| number of Cl- | 200 | 348 |
| total number of atoms | 485408 | 546884 |
| equilibrated box dimension (nm) | $9.7 \times 9.7 \times 39.8$ | $9.7 \times 9.7 \times 44.1$ |
| ionic strength (mM) | 106 | 107 |
| local equilibration benchmark | 46 ns/day* | 44 ns/day* |
| local equilibration length | 210 ns | 210 ns |
| Anton 2<sup>S12</sup> benchmark | 4.4 μs/machine day | 4.2 μs/machine day |
| Anton 2 equilibration | 0.15 μs NPT | 0.15 μs NPT |
| Anton 2 simulation length | 2 μs NVT | 2 μs NVT |

**Table S2.**
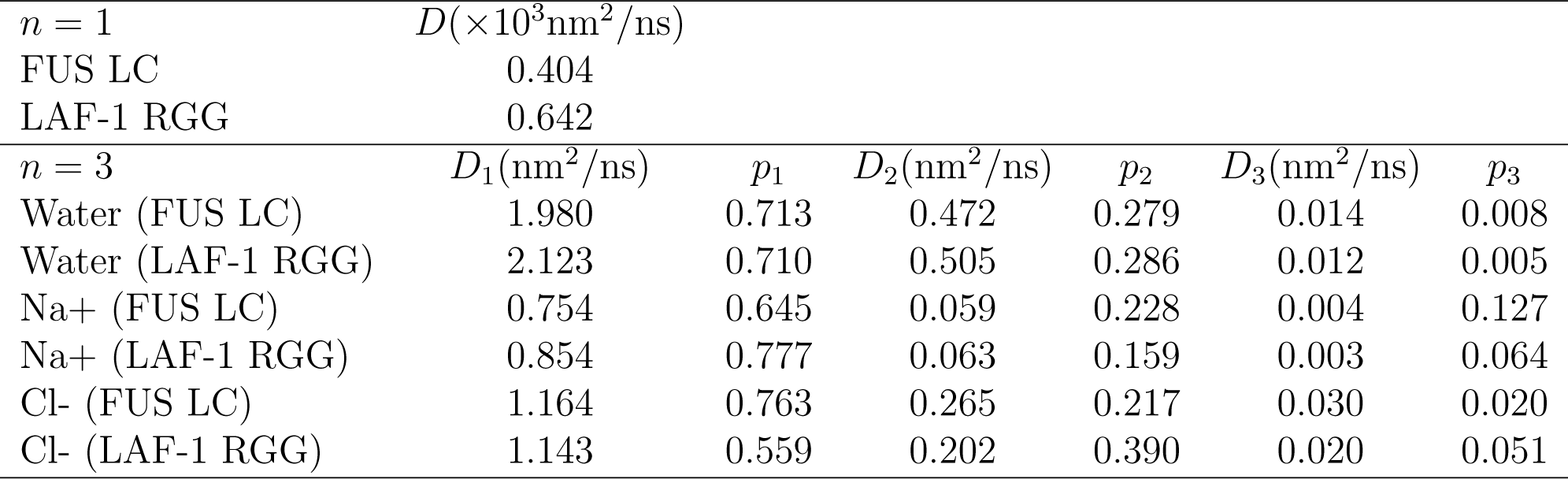
Diffusion coefficients of proteins, water and ions along *z*-axis. See supporting methods for details.

|  |  |  |  |  |  |  |
| --- | --- | --- | --- | --- | --- | --- |
| $n = 1$ | $D(\times 10^3 \text{nm}^2/\text{ns})$ | | | | | |
| FUS LC | 0.404 |  |  |  |  |  |
| LAF-1 RGG | 0.642 |  |  |  |  |  |
| $n = 3$ | $D_1(\text{nm}^2/\text{ns})$ | $p_1$ | $D_2(\text{nm}^2/\text{ns})$ | $p_2$ | $D_3(\text{nm}^2/\text{ns})$ | $p_3$ |
| Water (FUS LC) | 1.980 | 0.713 | 0.472 | 0.279 | 0.014 | 0.008 |
| Water (LAF-1 RGG) | 2.123 | 0.710 | 0.505 | 0.286 | 0.012 | 0.005 |
| Na+ (FUS LC) | 0.754 | 0.645 | 0.059 | 0.228 | 0.004 | 0.127 |
| Na+ (LAF-1 RGG) | 0.854 | 0.777 | 0.063 | 0.159 | 0.003 | 0.064 |
| Cl- (FUS LC) | 1.164 | 0.763 | 0.265 | 0.217 | 0.030 | 0.020 |
| Cl- (LAF-1 RGG) | 1.143 | 0.559 | 0.202 | 0.390 | 0.020 | 0.051 |

